# Reversion and non-reversion mechanisms of resistance to PARP inhibitor or platinum chemotherapy in *BRCA1/2*-mutant metastatic breast cancer

**DOI:** 10.1101/832717

**Authors:** Adrienne G. Waks, Ofir Cohen, Bose Kochupurakkal, Dewey Kim, Connor E. Dunn, Jorge Buendia Buendia, Seth Wander, Karla Helvie, Maxwell R. Lloyd, Lori Marini, Melissa E. Hughes, Samuel S. Freeman, S. Percy Ivy, Joseph Geradts, Steve Isakoff, Patricia LoRusso, Viktor A. Adalsteinsson, Sara M. Tolaney, Ursula Matulonis, Ian E. Krop, Alan D. D’Andrea, Eric P. Winer, Nancy U. Lin, Geoffrey I. Shapiro, Nikhil Wagle

**Affiliations:** Department of Medical Oncology, Dana-Farber Cancer Institute, Boston, MA; Department of Medicine, Brigham and Women’s Hospital, Boston, MA; Broad Institute of MIT and Harvard, Cambridge, MA; Harvard Medical School, Boston, MA; Center for Cancer Precision Medicine, Dana-Farber Cancer Institute, Boston, MA; Center for DNA Damage and Repair, Dana-Farber Cancer Institute, Boston, MA; Massachusetts General Hospital Cancer Center and Department of Medicine, Harvard Medical School, Boston, MA; Investigational Drug Branch, Cancer Therapy Evaluation Program, National Cancer Institute, Bethesda, MD; City of Hope Comprehensive Cancer Center, Duarte, CA; Yale Cancer Center, New Haven, CT; Department of Radiation Oncology, Dana-Farber Cancer Institute and Harvard Medical School; University of Massachusetts Medical School, Worcester, MA

**Keywords:** *BRCA1*, *BRCA2*, breast cancer, PARP inhibitor, platinum

## Abstract

**Background:** Little is known about mechanisms of resistance to PARP inhibitors and platinum chemotherapy in patients with metastatic breast cancer and *BRCA1*/2 mutations. Further investigation of resistance in clinical cohorts may point to strategies to prevent or overcome treatment failure.

**Patients and Methods:** We obtained tumor biopsies from metastatic breast cancer patients with *BRCA1/2* deficiency before and after acquired resistance to PARP inhibitor or platinum chemotherapy. Whole exome sequencing was performed on each tumor, germline DNA, and circulating tumor DNA. Tumors underwent RNA sequencing, and immunohistochemical staining for RAD51 foci on tumor sections was performed for functional assessment of intact homologous recombination.

**Results:** Pre- and post-resistance tumor samples were sequenced from 8 patients (4 with *BRCA1* and 4 with *BRCA2* mutation; 4 treated with PARP inhibitor and 4 with platinum). Following disease progression on DNA-damaging therapy, four patients (50%) acquired at least one somatic reversion alteration likely to result in functional BRCA1/2 protein detected by tumor or circulating tumor DNA sequencing. Two patients with germline *BRCA1* deficiency acquired genomic alterations anticipated to restore homologous recombination through increased DNA end resection: loss of *TP53BP1* in one patient and amplification of *MRE11A* in another. RAD51 foci were acquired post-resistance in all patients with genomic reversion, consistent with reconstitution of homologous recombination. All patients whose tumors demonstrated RAD51 foci post-resistance were intrinsically resistant to subsequent lines of DNA-damaging therapy.

**Conclusions:** Genomic reversion in *BRCA1/2* was the most commonly observed mechanism of resistance, occurring in 4 of 8 patients. Novel sequence alterations leading to increased DNA end resection were seen in two patients, and may be targetable for therapeutic benefit. The presence of RAD51 foci by immunohistochemistry was consistent with BRCA1/2 protein functional status from genomic data and predicted response to later DNA-damaging therapy, supporting RAD51 focus formation as a clinically useful biomarker.

## Introduction

Approximately 5% of breast cancer patients carry germline mutations in *BRCA1* or *BRCA2*,^1^ tumor suppressor genes that function in the repair of DNA double stranded breaks by homologous recombination (HR).^2–4^ Multiple lines of evidence from *in vitro* experiments to large-scale randomized clinical trials have demonstrated that BRCA1/2-deficient cancers are particularly sensitive to two classes of DNA-damaging therapies: platinum chemotherapy and poly(adenosine diphosphate-ribose) polymerase (PARP) inhibitors.^1,5,6^ Platinum agents are increasingly widely used in patients with metastatic breast cancer and germline *BRCA1/2* mutations, and two PARP inhibitors (PARPi; olaparib and talazoparib) obtained United States Food and Drug Administration (FDA) approval for this indication in 2018.^7^ As the clinical use of these agents expands, it is increasingly important to understand how resistance occurs and to develop biomarkers predictive of response.

Previously described mechanisms of resistance to PARPi or platinum chemotherapy fall into two main categories: alteration of a protein in the HR pathway (including acquired re-expression of functional BRCA protein, known as reversion),^8–10^ and altered expression of a protein in the replication fork protection pathway.^11,12^ Loss of drug efficacy related to overexpression of drug efflux pumps or desmoplastic stromal reaction in tumor tissue has also been reported.^10^ Though *BRCA1* or *BRCA2* reversions have been described in many clinical cohorts, non-reversion mechanisms of resistance are almost exclusively described in *in vitro* models, and there is great need to explore their relevance in clinical specimens. Furthermore, much of the clinical work to date has been in ovarian cancer and prostate cancer,^10,13–15^ with fewer investigations of PARPi or platinum resistance in breast cancer patients.^16,17^

The goal of this study was to use tumor sequencing to identify both reversion and non-reversion mechanisms of acquired resistance to PARPi or platinum chemotherapy in patients with *BRCA1/2*-deficient metastatic breast cancer. Sequencing of the whole exome and the whole transcriptome was performed in paired tumor specimens before and after acquired resistance to PARPi or platinum. Presence of RAD51 foci (a marker of intact HR) was also measured to assess the functional impact of alterations identified in sequencing results. Our findings suggest two novel non-reversion mechanisms of resistance, and support RAD51 focus formation as a clinically useful biomarker of resistance to PARPi and platinum chemotherapy.

## Methods

Further details can be found in Supplemental Methods

### Patient cohort and biopsies

Prior to any study procedures, all patients provided written informed consent for research biopsies and sequencing of tumor and normal DNA, as approved by the Dana-Farber/Harvard Cancer Center Institutional Review Board (DF/HCC Protocol 05-246). We identified all patients enrolled on this sequencing study who had germline or somatic deleterious *BRCA1* or *BRCA2* alteration and a tissue biopsy performed following acquired resistance to PARPi or platinum therapy between July 2015 to July 2017. Acquired resistance was defined as disease progression following either complete response, partial response, or stable disease for a period of >3 months, followed by disease progression on therapy. Patient/tumor characteristics and breast cancer treatment history, including response to each treatment received assessed by a breast medical oncologist, were extracted from the medical record.

### Tumor and blood sequencing

DNA extraction and construction of libraries for massively parallel sequencing were performed as previously described.^18^ Cell-free DNA was isolated and circulating tumor DNA was sequenced using the ichorCNA method, as previously described.^19^ Analysis pipeline details follow in Supplemental Methods.

### Identification of reversions in *BRCA1* and *BRCA2*

Short frame-restoring indels of length < 100bp were identified using the 2-out-of-3 voting scheme described in the Supplemental Methods regarding somatic alterations. Longer deletions of length >= 100bp were identified using SvABA, and the resulting VCF was annotated using svaba-annotate.R and AnnotSV version 1.2.^20^ Long deletions identified in *BRCA1* and *BRCA2* were checked for validity and impact to the reading frame via manual inspection of the raw reads (*.alignments.txt.gz) aligned to contigs assembled by SvABA.

### Immunohistochemical staining

For RAD51 staining, serial sections of FFPE tumor biopsies were stained as previously described^21^ using antibodies to RAD51 and Geminin independently. A sample was classified as HR proficient if more than three RAD51 foci were present in a minimum of one cell in three 40X fields. If RAD51 foci were absent, the sample was classified as HR deficient if greater than 3% of the cells were Geminin positive. If there were no RAD51 foci and less than 3% of the cells were Geminin positive, the proliferation rate of the tumor was classified as low and HR status could not be determined. FFPE sections of a cell line block containing irradiated and unirradiated HR-proficient (HCC1569) and HR-deficient (MDA-MB-436) breast cancer cell lines were used as positive and negative controls.^21^

For phosphorylated replication protein A (phospho-RPA) staining, FFPE tumor sections were stained using the standardized immunohistochemical protocol for phospho RPA32 (S4/S8) antibody A300-245A (Bentyl Laboratories). Stained slides were scanned and image analysis was performed on the Aperio platform (Leica Biosystems).

## Results

### Patient and tumor characteristics

Eight patients with metastatic breast cancer, germline and/or somatic inactivating mutation in *BRCA1* or *BRCA2*, and acquired resistance to any PARPi or platinum chemotherapy were identified (**Table 1**). Seven patients were germline mutation carriers and the eighth had a somatic mutation only. The majority of patients had hormone receptor-positive (HR+) tumors due to the fact that the initial focus of the umbrella collection protocol was on HR+ breast cancer. Four of eight patients had acquired resistance to PARPi, and four had acquired resistance to platinum chemotherapy.

**Table 1.** Clinico-pathologic and treatment data for the cohort. Anatomic stage is indicated. Best overall response was determined by Response Evaluation Criteria in Solid Tumors (if patient on a clinical trial) or chart review by a breast medical oncologist (if patient not on a clinical trial). *One patient (#303) had a somatic mutation in *BRCA2*. **This patient underwent blinded randomization to veliparib or placebo on a clinical trial. *Abbreviations: CR, complete response; dx, diagnosis; ER, estrogen receptor; fs, frameshift; ns, nonsense mutation; PD, progressive disease; PaR, partial response; PR, progesterone receptor; SD, stable disease; tx, treatment; UNK, unknown*

| Patient ID | Germline mutation* | Age at dx (yrs) | Stage at dx | Receptor status at dx | DNA-damaging tx received | Duration of response | Best overall response | Reason for stopping tx |
| --- | --- | --- | --- | --- | --- | --- | --- | --- |
| 292 | *BRCA1* (fs) | 44 | II | ER+/PR+/HER2- | Carboplatin | 9 mos | SD | PD |
| 303 | *BRCA2* (splice site, somatic) | 43 | III | ER+/PR+/HER2- | Cisplatin | 15 mos | CR | PD |
| 318 | *BRCA2* (fs) | 51 | I | ER+/PR+/HER2- | Olaparib | 11 mos | SD | PD |
| 339 | *BRCA2* (fs) | 30 | III | ER+/PR+/HER2- | Carboplatin | 6 mos | UNK | PD |
| 349 | *BRCA1* (ns) | 29 | I | ER+/PR+/HER2- | Olaparib | 11 mos | PaR | PD |
| 359 | *BRCA1* (fs) | 53 | II | ER+/PR-/HER2- | Olaparib | 4 mos | SD | PD |
| 510 | *BRCA2* (fs) | 58 | III | ER+/PR+/HER2- | Carboplatin +/- veliparib** | 9 mos | PaR | PD |
| 565 | *BRCA1* (fs) | 42 | II | ER-/PR-/HER2- | Veliparib | 20 mos | PaR | PD |

### Reversions identified in *BRCA1* and *BRCA2* following exposure to platinum chemotherapy or PARPi

Four of eight patients demonstrated definite or putative reversion to a functional *BRCA1* or *BRCA2* open reading frame following acquired resistance to PARPi or platinum chemotherapy. Details of reversion events for all four patients are shown in **Figure 1**. The identified reversions suggested restoration of intact HR through reconstitution of functional BRCA1/2 protein as a likely mechanism of resistance to PARPi/platinum chemotherapy in these four patients. Consistent with this, in all four patients the pre-resistance tumors had no RAD51 foci (indicative of defective HR) while the post-resistance tumors demonstrated the acquisition of RAD51 foci (indicative of restoration of HR) (**Figure 1A-D**). **Table S1** shows the exact genomic coordinates, allelic fraction, and cancer cell fraction for each reversion event. We categorized reversion events as definite (patients 318, 339, and 349) if acquired restoration of *BRCA1* or *BRCA2* open reading frame could be concluded directly from gene sequence for at least one genomic event, and as putative (patient 510) if open reading frame sequence could not be directly concluded but genomic events nearby to the original inactivating event were newly acquired following resistance to DNA-damaging therapy and accompanied by acquisition of RAD51 foci post-resistance.

**Figure 1.**
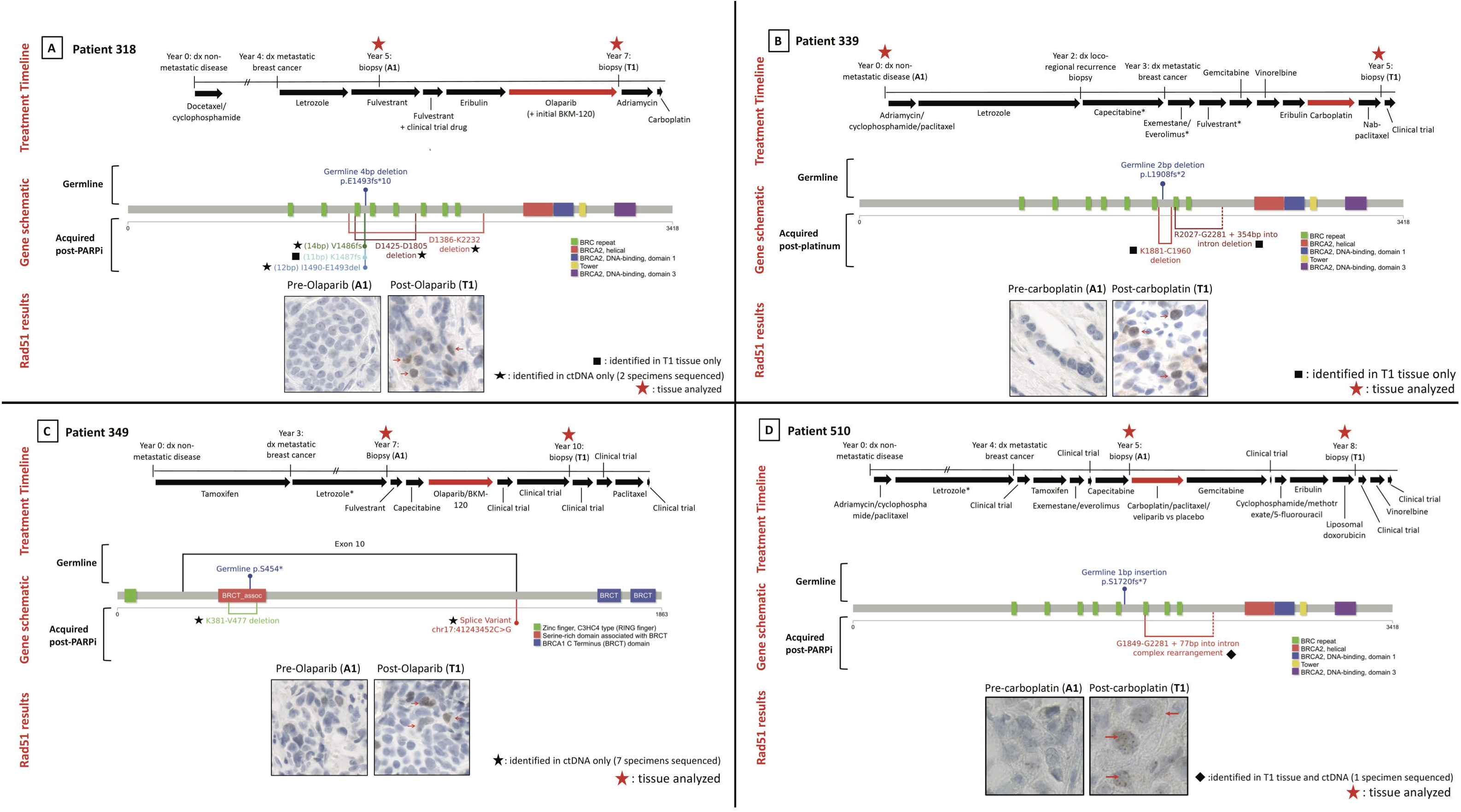
Reversions identified in *BRCA1* and *BRCA2* following exposure to PARP inhibitor or platinum. **(A)** Following olaparib exposure, patient 318 acquired three short deletions immediately upstream of or encompassing the germline frameshift deletion in *BRCA2*, and two long in-frame deletions encompassing the germline frameshift deletion and contained within *BRCA2* exon 11. Each of these acquired alterations is expected to restore *BRCA2* open reading frame, re-establishing HR proficiency, and this is supported by the reacquisition of RAD51 foci in the post-resistance biopsy. **(B)** Following exposure to carboplatin, patient 339 acquired an in-frame deletion encompassing the germline frameshift deletion and contained within *BRCA2* exon 11, with accompanying reacquisition of RAD51 foci. This patient also acquired a somatic deletion just downstream of the germline frameshift, reaching into the adjacent exon. **(C)** Patient 349 had a germline nonsense mutation in *BRCA1* exon 10 and acquired an exon 10 splice site mutation following resistance to olaparib, with accompanying reacquisition of RAD51 foci. In addition, this patient acquired an in-frame deletion encompassing the germline frameshift deletion. **(D)** Patient 510 had a germline frameshift insertion in *BRCA2* exon 11 and acquired a somatic deletion close downstream, beginning within exon 11 and reaching into the adjacent intron, after becoming resistant to carboplatin with or without veliparib. Figure notes: no clinical trials involving PARP inhibitor/platinum or other therapy specifically for *BRCA1/2*-mutant pts occurred between indicated sequenced biopsies. Treatment timelines are to scale unless noted; double hash marks indicate treatment duration longer than diagrammed. Red arrows identify cells with positive staining for RAD51 foci. *Exact treatment duration unknown. *Abbreviations: dx, diagnosis*

In a subset of patients, circulating tumor DNA (ctDNA) was available for sequencing in addition to tumor in a subset of patients. **Table S2** shows the number of circulating tumor DNA (ctDNA) specimens that successfully underwent WES from blood in each patient (median 2, range 0-7); 7 patients had successful sequencing of at least one ctDNA specimen (including all 4 patients with reversion events). All ctDNA specimens were drawn close in time or subsequent to post-resistance tumor sampling. In the 4 patients with reversion events, there were 10 total reversion events identified. Six events were found in blood only, 3 were found in tumor only, and 1 was found in blood and tumor, confirmed by manual review of unfiltered sequencing results from both tumor and blood for all events. Of note, two patients had reversions identified only in ctDNA; in both cases, ctDNA was sampled more than tumor.

### Genomic analysis of acquired resistance pathways identifies TP53BP1 loss and MRE11A amplification in two patients with *BRCA1* germline mutation

We compared tumor whole exome sequencing before and after acquired resistance to PARPi or platinum chemotherapy to identify potential non-reversion mechanisms of resistance to therapy. We focused our initial analysis on single nucleotide variants (SNVs) and copy number variants (CNVs) in 20 genes from pathways previously linked to PARPi/platinum response or resistance in preclinical models and/or clinical specimens.^9–12,16,22–30^ Genomic alterations involving these 20 genes in pre-resistance versus post-resistance tumor specimens from each patient are shown in **Figure 2**.

**Figure 2.**
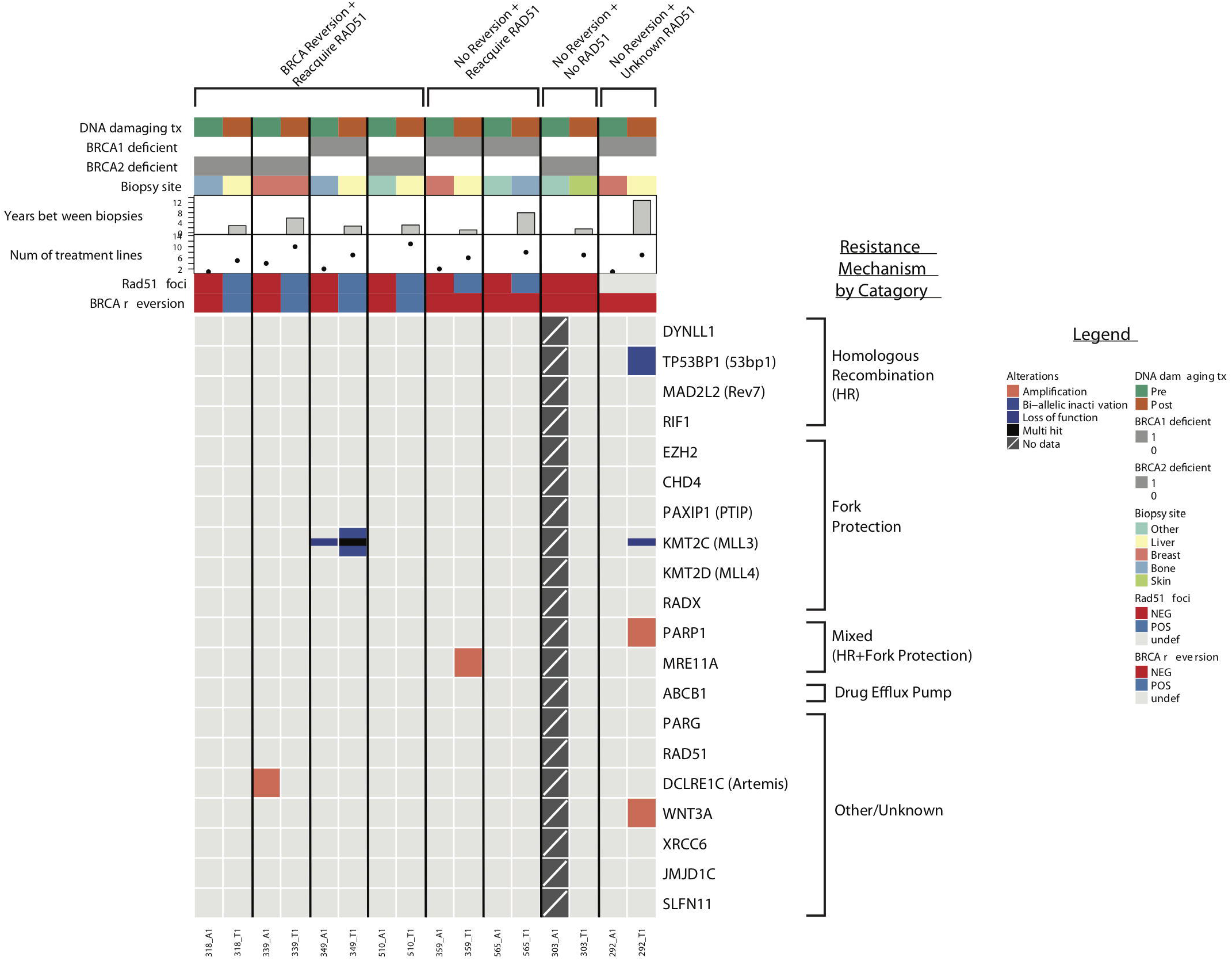
SNV and CNV events in 20 genes from PARPi/platinum resistance pathways. Co-mutation plot showing SNV and CNV events in 20 genes from pathways previously implicated in resistance to PARPi or platinum, across 15 metastatic breast cancer tumor samples obtained either before or after the acquisition of resistance to PARPi or platinum. Each column of data represents a unique tumor specimen; each row represents a gene of interest. No pre-resistance tumor sample was available for sequencing in patient 303. Horizontal tracks along the top of the plot indicate select clinical parameters for each specimen, presence or absence of detected *BRCA* reversion in either tumor tissue or ctDNA, and presence or absence of RAD51 foci staining.

In two patients, we identified acquired genomic alterations anticipated to lead to restoration of HR through increased DNA end resection. Patient 292, with a pathogenic germline mutation in *BRCA1* and no identified reversion in *BRCA1* after carboplatin treatment (**Figure 3A**), acquired bi-allelic inactivation in *TP53BP1* resulting from an antisense fusion between *TP53BP1* and *GALNT2* plus loss of heterozygosity (**Figure 3B**). Low expression of *TP53BP1* was also observed in the post-resistance tumor specimen. Loss of *TP53BP1* is expected to facilitate BRCA1-independent end resection following double stranded DNA breaks, since loss of 53BP1 *in vitro* restores HR in cells lacking BRCA1, leading to PARPi/platinum resistance despite maintained BRCA1 deficiency.^28,31,32^ Tissue was insufficient for RAD51 staining or 53BP1 protein staining in this patient.

**Figure 3.**
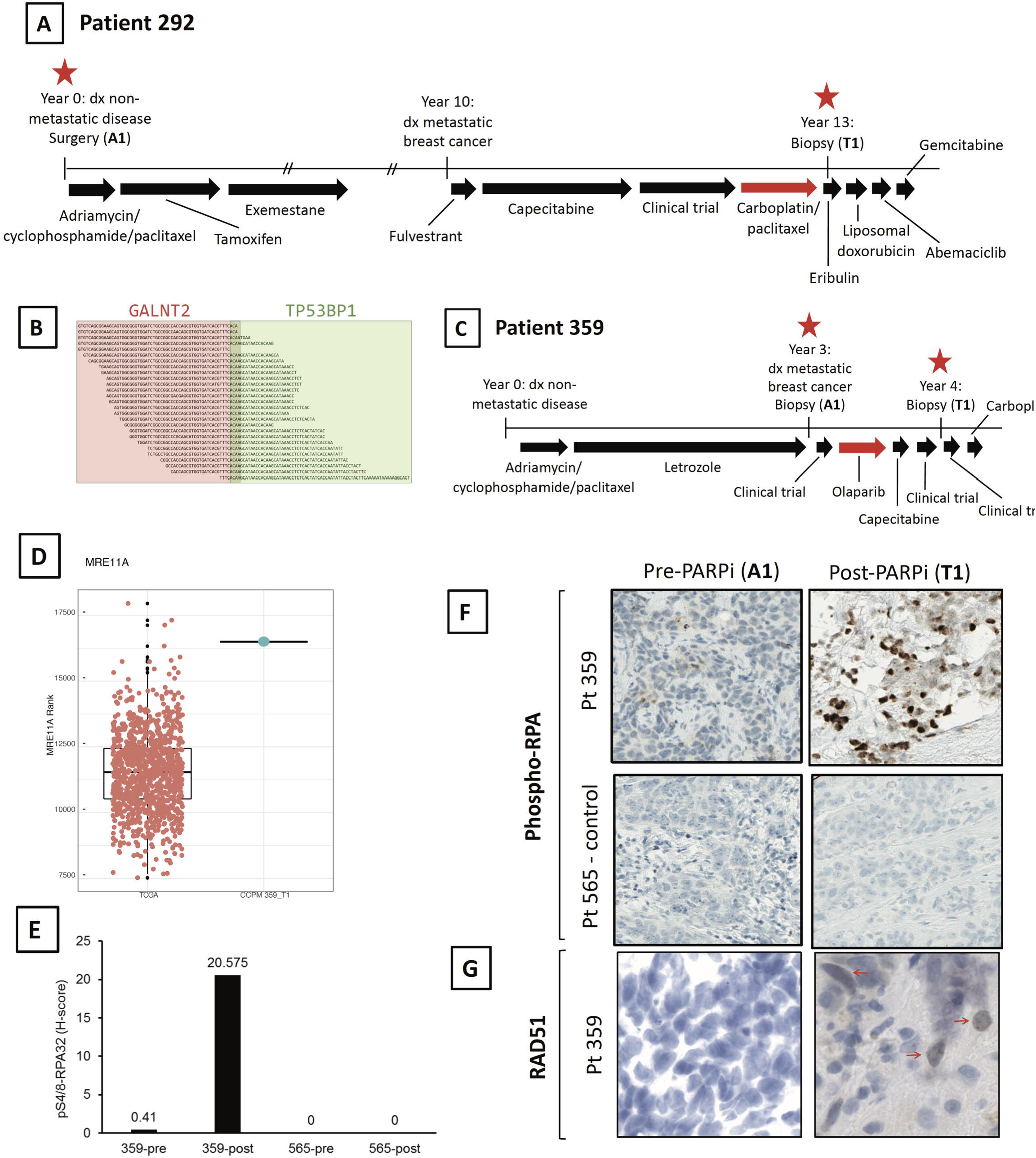
Genomic alterations in *TP53BP1* and *MRE11A* acquired in two patients with germline *BRCA1* deficiency. **(A)** Patient 292 had a germline deleterious *BRCA1* mutation and acquired resistance to a carboplatin-containing regimen. **(B)** In a post-resistance tissue biopsy, patient 292 acquired biallelic inactivation of the gene *TP53BP1* (loss of heterozygosity plus antisense fusion between *TP53BP1* and *GALNT2*). **(C)** Patient 359 had a germline deleterious *BRCA1* mutation and acquired resistance to olaparib. **(D)** In a post-resistance tissue biopsy, patient 359 showed very high RNA expression of *MRE11A*; figure shows comparison of *MRE11A* expression in all breast tumors from The Cancer Genome Atlas (N=1093; pink dots) versus *MRE11A* expression in patient 359 post-resistance specimen (blue dot; at 99.7^th^ percentile of TCGA samples). **(E)** Pre- and post-resistance tumor biopsies from patient 359 show increase in phospho-RPA protein staining post-resistance; patient 565 (whose tumor showed no genomic evidence of acquired increased end resection post-resistance) is shown as a control. **(F)** Representative histology images of phospho-RPA stain for patient 359 pre- and post-resistance (patient 565 is again shown as a control). **(G)** Reacquisition of RAD51 foci following acquired resistance to olaparib in patient 359. Red arrows identify cells with positive staining for RAD51 foci. Treatment timelines are to scale unless noted; double hash marks indicate treatment duration longer than diagrammed. *Abbreviations: dx, diagnosis*

Patient 359, who had a pathogenic germline mutation in *BRCA1* and no identified reversion in *BRCA1* following exposure to olaparib (**Figure 3C**), acquired amplification of *MRE11A*, which encodes a DNA exonuclease that functions in end resection.^30^ *MRE11A* amplification could plausibly lead to PARPi resistance by increasing end resection at double stranded DNA breaks, therefore restoring HR proficiency in tumor cells despite *BRCA1* deficiency. Supporting the functional relevance of the *MRE11A* amplification in causing resistance to PARP inhibition in this patient, RNA expression of MRE11A was high in the post-resistance tumor specimen (**Figure 3D**), and tumor staining for phospho-RPA (a marker of DNA end resection) was substantially increased post-resistance (**Figure 3E-F**). Moreover, RAD51 foci were re-acquired at the post-olaparib timepoint (**Figure 3G**), as would be expected for a tumor with HR restoration through increased end resection. In this sample that increased DNA end resection and restored HR without *BRCA1* somatic reversion, we propose that the acquisition of *MRE11A* amplification is the likely biological mechanism of olaparib resistance.

One additional acquired genomic alteration that could potentially contribute to therapeutic resistance was seen in the 20 genes analyzed. Patient 349, with a pathogenic germline mutation in *BRCA1* and a putative somatic reversion in *BRCA1*, acquired biallelic inactivation of *KMT2C*, a histone methyltransferase necessary for the presence of MRE11 at replication forks, among many other cellular functions.^12,33^ The loss of *KMT2C* in this patient suggests that a replication fork-stabilizing event may have occurred that subsequently conferred PARPi resistance.

Broader analysis of genomic alterations occurring in a larger set of 276 genes shown to be involved in all DNA damage repair processes in cancer^34^ did not reveal any explanatory mechanism of resistance beyond those identified above. In addition, given previous evidence indicating that gene fusions driving overexpression of *Abcb1* (drug efflux pump) can cause resistance to PARPi,^10,16^ all gene fusion events were examined in the cohort. No fusions in *Abcb1* or other genes of interest were identified. Following all analyses, we did not identify any candidate mechanisms in patients 303 or 565, whose acquired resistance remained unexplained.

### RAD51 foci and resistance to subsequent lines of platinum chemotherapy or PARPi

Of the six patients who acquired RAD51 foci following exposure to platinum chemotherapy or PARPi, three went on to have subsequent exposure to a different DNA-damaging therapy in a later line of treatment (**Table 2**). In all three cases, patients displayed intrinsic resistance to subsequent PARPi and/or platinum. By contrast, RAD51 foci were absent from all tested tumors prior to initial therapy with PARPi or platinum (**Figure 2**), among patients selected for initial response to therapy. The data are consistent with the premise that intact or impaired HR, measured by the presence or absence of RAD51 foci, correlates with response to PARPi/platinum agents in *BRCA1/2*-deficient tumors, though the number of patients in this cohort is too small to allow definitive conclusions about the clinical utility of RAD51 focus formation as a predictive biomarker.

**Table 2.** Reversion status and RAD51 status following platinum/PARP inhibitor exposure, and clinical response to subsequent platinum/PARP inhibitor. Intrinsic resistance is defined as progressive disease at first restaging or clinical progression prior to first restaging. *Abbreviations: NA, not applicable; tx, treatment*

| Patient ID | BRCA reversion status post-DNA damaging tx | RAD51 foci status pre-DNA damaging tx | RAD51 foci status post-DNA damaging tx | Subsequent DNA damaging tx exposure | Best response to subsequent DNA damaging tx |
| --- | --- | --- | --- | --- | --- |
| 292 | No | Unknown | Unknown | None | NA |
| 303 | No | Absent | Absent | None | NA |
| 318 | Yes, definite | Absent | **Present** | **Carboplatin** | **Intrinsic resistance** |
| 339 | Yes, definite | Absent | **Present** | None | NA |
| 349 | Yes, definite | Absent | **Present** | None | NA |
| 359 | No | Absent | **Present** | **Carboplatin** | **Intrinsic resistance** |
| 510 | Yes, putative | Absent | **Present** | None | NA |
| 565 | No | Absent | **Present** | **Carboplatin and olaparib (as separate regimens)** | **Intrinsic resistance to both** |

## Discussion

In this study, we used whole exome sequencing of tumor and blood, RNA sequencing of tumor, and formation of RAD51 foci by immunohistochemistry to interrogate resistance to PARP inhibitors or platinum chemotherapy and its correlation with tumor HR proficiency in a cohort of patients with metastatic breast cancer. While the cohort is only eight patients, to our knowledge it is the largest such investigation looking at mechanisms of resistance in *BRCA1/2*-deficient tumor tissue from metastatic breast cancer patients. Reversions were identified in one-half of patients. Moreover, sequencing data suggest additional, biologically plausible non-reversion mechanisms of resistance as well, including amplification of *MRE11A* and biallelic inactivation of *TP53BP1*, each of which may restore HR through increased DNA end resection. Lastly, the presence or absence of RAD51 foci correlated with resistance or response, respectively, to DNA-damaging therapy, implying that immunohistochemical assessment of RAD51 should be further explored as a clinically deployable predictive biomarker to guide therapeutic decisions in *BRCA1/2*-deficient tumors.

Reversion to protein-coding *BRCA1* or *BRCA2* transcript has been previously reported following exposure to platinum chemotherapy and/or PARPi both in preclinical models of *BRCA1/2*-deficient tumor cells and in patients with *BRCA1/2*-mutated breast cancer, ovarian cancer, and prostate cancer.^10,13,14,17,35,36^ The reversions identified in this cohort occurred through a variety of genomic mechanisms, including short deletions nearby to or encompassing a germline frameshift alteration (patient 318), long deletions encompassing a germline frameshift alteration (patients 318, 339, and 349), and deletions or splice site mutations postulated to activate alternative splicing to a functional though possibly hypomorphic BRCA1/2 protein (patients 349 and 510). Prior evidence supports the biological plausibility of each of these revertant mechanisms: BRCA2 proteins with large amounts of protein-coding sequence deleted have been demonstrated following PARPi exposure, and retain HR functionality;^36,37^ and activation of alternative splice isoforms has been associated with PARPi or platinum resistance in both *BRCA1*- and *BRCA2*-deficient tissue.^26,35,38^ Our similar observations support the clinical relevance of these events.

It is interesting that despite co-sampling of tissue and ctDNA in 7 of 8 patients, and identification of reversions using both methods, the majority of reversion events were not shared between tumor and blood specimens. The preponderance of events identified in blood only may reflect the fact that multiple post-resistance blood specimens were sequenced in most patients (compared to only a single post-resistance tumor specimen sequenced in all patients). The discordance also highlights the limitations of a single tumor or blood sample in isolation to comprehensively capture heterogeneous genomics across multiple different metastatic lesions at distinct timepoints, and is consistent with a previous report showing incomplete overlap between reversions identified in blood versus tumor.^13^

Though genomic reversion is a frequently reported mechanism of clinical resistance to PARPi or platinums among patients with *BRCA1/2*-mutant tumors, as observed in our cohort, not all patients revert. At present there are no known parameters to predict which patients will acquire somatic reversions and which will not, though this information could carry immense clinical utility as it may identify individuals who would benefit from upfront combined therapeutic strategies to prevent development of resistance. Interestingly, each of the four patients we observed to acquire somatic reversions harbored their original germline mutation within the longest exon of *BRCA1/2* transcript (exons 10 and 11 for *BRCA1* and *BRCA2*, respectively). These exons are both above the 99^th^ percentile for exon length across the human genome), suggesting that perhaps replicative instability of very long exons—particularly with respect to long deletions—could predispose a patient to develop reversions. In addition, a very similar combination of germline mutation and somatic reversion as observed in patient 318 has been previously reported in a patient with ovarian cancer and *BRCA2* mutation.^39^ Together these observations raise the question of whether the location and/or type of germline mutation in *BRCA1/2* may assist in predicting which patients will experience reversion and which will not. Work to compile and map reversions and associated germline mutations identified to date across tumor types is warranted to address this question.

Non-reversion mechanisms of resistance to PARPi/platinum have been uncommonly identified in clinical specimens; we identified two non-revertant patients in whom genomic evidence supports the acquisition of resistance through upregulation of DNA end resection. Loss of 53BP1, a protein involved in DNA end resection, has been shown to restore HR functionality in *BRCA1*-deficient cells, and to eliminate the cells’ platinum/PARPi sensitivity.^31,32^ Reduced 53BP1 has also been described in platinum and PARPi-resistant ovarian cancer patient tumor specimens and patient-derived xenograft models but to our knowledge has never been demonstrated as a mechanism of resistance in breast tumor specimens.^25,40,41^ 53BP1 normally inhibits the activity of MRE11, a DNA exonuclease, at DNA double stranded breaks.^12^ *MRE11A* amplification (seen here in patient 359 post-olaparib) has not previously been reported as a mechanism of resistance to PARPi/platinum, but is entirely plausible as an alternative means to promote DNA end resection. This biology is supported by increased phospho-RPA staining in our patient’s tumor, and represents a potential novel mechanism of resistance identified in this cohort.

It should be noted that MRE11 also plays a role in replication fork degradation, and in this context a theoretical consequence of its amplification could actually be increased sensitivity to PARPi/platinum (i.e. opposite directionality from its effect on double stranded break repair).^12,28^ However, the broad evidence supporting 53BP1 loss as a resistance mechanism, the increase in phospho-RPA staining, and the fact that this patient’s tumor regained RAD51 foci, all contradict this as a predominant biological effect in this tumor. Conversely, in the case of patient 349 with acquired biallelic inactivation of *KMT2C*, *KMT2C* loss could have prevented MRE11 access to replication forks, creating a fork stabilizing event contributing to PARPi resistance. It is noteworthy that this event occurred with a concomitant putative *BRCA1* reversion. In ovarian cancer cell line models of PARPi resistance, restoration of HR and stabilization of replication forks have indeed been shown to occur together,^28^ suggesting that BRCA1-deficient cells may employ multiple strategies to overcome the pressure of continued PARPi exposure.

Overall, our results are intriguing as they represent the first direct evidence of increased DNA end resection via 53BP1 loss or MRE11 upregulation as a clinically relevant mechanism of resistance to PARPi/platinum in *BRCA1*-deficient breast tumors. Though numbers are too small to draw any conclusions about the broader prevalence of these mechanisms, it is notable that two of four patients with *BRCA1* mutation in our cohort acquire resistance in this manner. Examination of the DNA end resection pathway—53BP1 and MRE11 in particular—is warranted in larger cohorts of patients with *BRCA1* mutation and PARPi/platinum resistance.

Our results suggest that immunohistochemical staining for RAD51 foci offers real-time assessment of a tumor’s HR proficiency, and correlates with response and resistance to PARPi/platinum therapy. The feasibility of RAD51 foci staining in human tumors without antecedent exposure to DNA damaging agents has been previously demonstrated in two small cohorts,^25,42^ and our results offer further proof-of-concept. The presence of RAD51 foci has been shown to correlate with decreased efficacy of PARP inhibition in patient derived xenografts and in a cohort of 8 patients with germline *BRCA1/2* mutation.^25,42^ However, we demonstrate here for the first time that the presence or absence of RAD51 staining changes over time as predicted with HR-restoring mechanisms of resistance. Taken together, our data along with previous results strongly suggest that RAD51 staining should be investigated as a predictive biomarker in larger cohorts of *BRCA1/2*-deficient patients treated with PARPi/platinum.

Our study has several limitations. The cohort size is small, though the availability of paired tumor tissue from 7 of 8 patients offers a unique opportunity to closely examine resistance mechanisms, and to our knowledge this represents the largest such cohort specifically of metastatic breast cancer patients reported to date. As this was not a treatment-based clinical trial, therapies received and biopsy timepoints are heterogeneous, and the biopsies performed do not exactly bracket the PARPi/platinum treatments received. While it is therefore not possible to conclude that the resistance mechanisms identified specifically resulted from selective pressure of PARPi/platinum, each mechanism highlighted has been previously reported to result from PARPi/platinum exposure, or, in the case of *MRE11A* amplification, is in a known resistance pathway.

In conclusion, we report on a cohort of eight patients with metastatic breast cancer and *BRCA1/2*-deficient tumors who acquired resistance to PARPi or platinum therapy. Using whole exome and RNA sequencing, with supportive functional evidence from RAD51 foci staining in each case, we identified biologically plausible mechanisms of resistance in 6 patients: four with acquired somatic reversion in *BRCA1* or *BRCA2*, and two with acquired upregulation of DNA end resection. The fact that HR restoration explained resistance in the majority of this cohort suggests that HR-disrupting strategies (such as inhibition of phosphoinositide 3-kinase or cyclin-dependent kinases), or strategies disrupting both HR and replication fork stability (such as inhibition of ATR or CHK1)^9^ may represent the best opportunities to re-sensitize patients to PARPi or platinum therapies.

## Supporting information

Extended References

## Notes

**Funding:** Supported by an American Society of Clinical Oncology Young Investigator Award (AGW), the Breast Cancer Research Foundation (AGW, NUL), Specialized Program of Research Excellence (SPORE) in Breast Cancer NIH grant P50 CA168504 (AGW, IEK, ADD, EPW, NUL, GIS, NW), an NCI-Cancer Therapy Evaluation Program (CTEP) Biomarker Supplement to NIH grant UM1 CA186709 (BK, ADD, GIS), the Fashion Footwear Association of New York (to Dana-Farber Cancer Institute Breast Oncology Program), the National Comprehensive Cancer Network/Pfizer Collaborative Grant Program (NUL), Friends of Dana-Farber Cancer Institute (to N.U.L.), Yale Cancer Center grant 1UM1CA86689-05 (PL), Department of Defense W81XWH-13-1-0032 (N.W.), AACR Landon Foundation 13-60-27- WAGL (N.W.), Susan G. Komen CCR15333343 (N.W.), The V Foundation (N.W.), The Breast Cancer Alliance (N.W.), The Cancer Couch Foundation (N.W.), Twisted Pink (N.W.), Hope Scarves (N.W.), ACT NOW (to Dana-Farber Cancer Institute Breast Oncology Program). In addition, we thank Dana-Farber/Harvard Cancer Center in Boston, MA, for the use of the Specialized Histopathology Core, which provided immunohistochemical staining. Dana-Farber/Harvard Cancer Center is supported in part by an NCI Cancer Center Support Grant # NIH 5 P30 CA06516.

**Conflicts of Interest:** A.W. receives institutional research funding from Genentech and MacroGenics. S.W. reports consulting/advisory board role for Foundation Medicine, InfiniteMD, Eli Lilly, and Puma Biotechnology; and equity in InfiniteMD. S.S.F. filed a patent (WO2017161175A1) on methods applied in this study. S.I. receives research funding support from Genentech, PharmaMar, AstraZeneca, Merck (all to institution); and consulting fees from Genentech, Hengrui, Puma, Immunomedics, and Myriad. P.L reports serving on advisory boards at AbbVie (2018-2019), Alexion (2016-2017), Ariad (2016-2017), GenMab (2016-2018), Glenmark (2016-2017), Menarini (2016-2017), Novartis (2016-2017), CytomX (2016-2019), Omniox (2016-2017), Ignyta (2016-2017), Genentech (2016-2019), Takeda (2017-2020), SOTIO Consultant (2018-2019), Cybrexa (2018-2019), Agenus (2018-2020), IQVIA (2019-2020), TRIGR (2019-2020), Pfizer (2019-2020), I-MAB (2019-2020), ImmunoMet (2018-2020), Black Diamond (2019-2020), Sartarius (2019-2020), and Glaxo-Smith Kline (2019-2020); data safety monitoring boards/committees at Agios (2016-2019), Five Prime (2017-2020), Halozyme (2016-2019), FivePrime (2017-2019), and Tyme (2018-2020); and reports work with the imCORE Alliance at Roche-Genentech (2016-2019). V.A.A. filed a patent (WO2017161175A1) on methods applied in this study. S.M.T reports receiving institutional research support from Merck, Bristol-Myers Squibb, Exelixis, Eli Lilly, Pfizer, Novartis, AstraZeneca, Eisai, Nektar, Odenate, Sanofi, and Genentech and has served on advisory boards for Genentech, Eli Lilly, Novartis, Pfizer, Nektar, Immunomedics, Nanostring, Daiichi-Sankyo, Bristol-Meyers Squibb, Sanofi, Athenex, AstraZeneca, Eisai, Puma, and Merck. U.M. reports serving on advisory boards (paid) at Astrazeneca, Myriad Genetics, Clovis, Eli Lilly, Mersana, Geneos, Fuji Film, Cerulean; and consulting (paid) for Merck, 2X Oncology and Immunogen. I.E.K. receives institutional research funding from Genentech/Roche, Pfizer, Daiichi-Sankyo; has advisory board (honoraria) roles at Genentech/Roche, Daiichi-Sankyo, Macrogenics, Context Therapeutics, Taiho Oncology; and reports DSMC (honoraria) at Merck; and reports DSMB (honoraria) at Novartis. E.P.W. institutional research funding from Genentech/Roche and Merck; consultant/honoraria from Carrick Therapeutics, Genentech/Roche, Genomic Health, GSK, Jounce, Lilly, Merck, Seattle Genetics; advisory board/honoraria from Leap. A.D.D. reports receiving commercial research grants from Eli Lilly & Company, Sierra Oncology, and EMD Serono and is a consultant/advisory board member for Eli Lilly & Company, Sierra Oncology, and EMD Serono. G.I.S. has received research funding from Eli Lilly, Merck KGaA/EMD-Serono, Merck, and Sierra Oncology. He has served on advisory boards for Pfizer, Eli Lilly, G1 Therapeutics, Roche, Merck KGaA/EMD-Serono, Sierra Oncology, Bicycle Therapeutics, Fusion Pharmaceuticals, Cybrexa Therapeutics, Astex, Almac, Ipsen, Bayer, Angiex, and Daiichi Sankyo. N.U.L. reports institutional research funding from Genentech, Merck, Pfizer, Seattle Genetics; and consulting/ad board roles at Puma, Daichii, Seattle Genetics. N.W. was previously a stockholder and consultant for Foundation Medicine; has been a consultant/advisor for Novartis and Eli Lilly; and has received sponsored research support from Novartis and Puma Biotechnology. None of these entities had any role in the conceptualization, design, data collection, analysis, decision to publish, or preparation of the manuscript. All other authors report no conflicts of interest.

