## Supplemental Methods for "Reversion and non-reversion mechanisms of resistance to PARP inhibitor or platinum chemotherapy in *BRCA1/2*-mutant metastatic breast cancer"

*Patient cohort and biopsies*

Metastatic core biopsies were obtained from patients and samples were immediately snap frozen in OCT and stored in -80°C. Archival formalin-fixed paraffin-embedded (FFPE) tumor tissue from a pre-resistance timepoint was also obtained if available for each patient. A blood sample was obtained during the course of treatment, and whole blood was stored at -80°C until DNA extraction from peripheral blood mononuclear cells (for germline DNA) was performed. Cell free DNA was obtained from plasma for circulating tumor DNA analysis.

*Whole exome sequencing*

DNA extraction: DNA extraction was performed as previously described.^1^ For whole blood, DNA is extracted using magnetic bead-based chemistry in conjunction with the Chemagic MSM I instrument manufactured by Perkin Elmer. Following red blood cell lysis, magnetic beads bind to the DNA and are removed from solution using electromagnetized rods. Several wash steps follow to eliminate cell debris and protein residue from DNA bound to the magnetic beads. DNA is then eluted in TE buffer. For frozen tumor tissue, DNA and RNA are extracted simultaneously from a single frozen tissue or cell pellet sample using the AllPrep DNA/RNA kit (Qiagen).  For FFPE tumor tissues, DNA and RNA are extracted simultaneously using Qiagen’s AllPrep DNA/RNA FFPE kit. All DNA is quantified using Picogreen.

Library Construction: DNA libraries for massively parallel sequencing were generated as previously described,^1^ with the following modifications: the initial genomic DNA input into the shearing step was reduced from 3µg to 10-100ng in 50µL of solution. For adapter ligation, Illumina paired-end adapters were replaced with palindromic forked adapters (purchased from Integrated DNA Technologies) with unique dual indexed 8 base index molecular barcode sequences included in the adapter sequence to facilitate downstream pooling. With the exception of the palindromic forked adapters, all reagents used for end repair, A-base addition, adapter ligation, and library enrichment PCR were purchased from KAPA Biosciences in 96-reaction kits. In addition, during the post-enrichment solid phase reversible immobilization (SPRI) bead cleanup, elution volume was reduced to 30µL to maximize library concentration, and a vortexing step was added to maximize the amount of template eluted.

Solution-phase hybrid selection: After library construction, hybridization and capture were performed using the relevant components of Illumina’s Rapid Capture Exome Kit and following the manufacturer’s suggested protocol, with the following exceptions: first, all libraries within a library construction plate were pooled prior to hybridization. Second, the Midi plate from Illumina’s Rapid Capture Exome kit was replaced with a skirted PCR plate to facilitate automation. All hybridization and capture steps were automated on the Agilent Bravo liquid handling system.

Preparation of libraries for cluster amplification and sequencing: After post-capture enrichment, library pools were then quantified using quantitative PCR (KAPA Biosystems) with probes specific to the ends of the adapters; this assay was automated using Agilent’s Bravo liquid handling platform. Based on qPCR quantification, libraries were normalized and denatured using 0.1 N NaOH on the Hamilton Starlet.

Cluster amplification and sequencing: Cluster amplification of denatured templates was performed according to the manufacturer’s protocol (Illumina) using HiSeq 2500 Rapid Run v1/v2, HiSeq 2500 High Output v4 or HiSeq 4000 v1 cluster chemistry and HiSeq 2500 (Rapid or High Output) or HiSeq 4000 flowcells. Flowcells were sequenced on HiSeq 2500 using v1 (Rapid Run flowcells) or v4 (High Output flowcells) Sequencing-by-Synthesis chemistry or v1 Sequencing-by-Synthesis chemistry for HiSeq 4000 flowcells. The flowcells were then analyzed using RTA v.1.18.64 or later. Each pool of whole exome libraries was run on paired 76np runs, with a two 8 base index sequencing reads to identify molecular indices, across the number of lanes needed to meet coverage for all libraries in the pool.

Sequence data processing: Exome sequence data processing was performed using established analytical pipelines at the Broad Institute. A BAM file was produced with the Picard pipeline (http://picard.sourceforge.net/), which aligns the tumor and normal sequences to the hg19 human genome build using Illumina sequencing reads. The BAM was uploaded into the Firehose pipeline (http://www.broadinstitute.org/cancer/cga/Firehose), which manages input and output files to be executed by GenePattern.^2^

Sequencing quality control: Quality control modules within Firehose were applied to all sequencing data for comparison of the origin for tumor and normal genotypes and to assess fingerprinting concordance. Cross-contamination of samples was estimated using ContEst.^3^

*Somatic alteration assessment*

MuTect^4^ was applied to identify somatic single-nucleotide variants. Indelocator (<http://www.broadinstitute.org/cancer/cga/indelocator)>, Strelka^5^, and MuTect2 (<https://software.broadinstitute.org/gatk/documentation/tooldocs/current/org_broadinstitute_gatk_tools_walkers_cancer_m2_MuTect2)> were applied to identify small insertions or deletions. A voting scheme with inferred indels requiring at least 2 out of 3 algorithms.

Artifacts introduced by DNA oxidation (so called OxoG) during sequencing were computationally removed using a filter-based method ^6^. In the analysis of primary tumors that are formalin-fixed, paraffin-embedded samples [FFPE] we further applied a filter to remove FFPE-related artifacts.^7^

Reads around mutated sites were realigned with Novoalign (www.novocraft.com/products/novoalign/) to filter out false positive that are due to regions of low reliability in the reads alignment. At the last step, we filtered mutations that are present in a comprehensive WES panel of 8,334 normal samples (using the Agilent technology for WES capture) aiming to filter either germline sites or recurrent artifactual sites. We further used a smaller WES panel of normal 355 normal samples that are based on Illumina technology for WES capture, and another panel of 140 normals sequenced without our cohort^8^ to further capture possible batch-specific artifacts. Annotation of identified variants was done using Oncotator^9^ (http://www.broadinstitute.org/cancer/cga/oncotator).

*Copy number and copy ratio analysis*

To infer somatic copy number from WES, we used ReCapSeg (http:// gatkforums.broadinstitute.org/categories/recapseg-documentation), calculating proportional coverage for each target region (i.e., reads in the target/total reads) followed by segment normalization using the median coverage in a panel of normal samples. The resulting copy ratios were segmented using the circular binary segmentation algorithm.^10^

To infer allele-specific copy ratios, we mapped all germline heterozygous sites in the germline normal sample using GATK Haplotype Caller^11^ and then evaluated the read counts at the germline heterozygous sites in order to assess the copy profile of each homologous chromosome. The allele-specific copy profiles were segmented to produce allele specific copy ratios.

*Cancer cell fraction and evolutionary analysis*

Analysis using ABSOLUTE: To properly compare SNVs and indels in paired metastatic and primary samples, we considered the union of all mutations called in either of the two samples. In order to re-evaluate the mutations that were not initially called, we used “forced calling” to quantify the number of reference and alternate reads at each mutations. These “force called” mutations in matched samples were used as input for ABSOLUTE^12^. The ABSOLUTE algorithm is using mutation-specific variant allele fractions (VAF) together with the computed purity, ploidy, and segment-specific allelic copy-ratio to compute cancer cell fractions (CCFs).

Analysis of evolution and clonal dynamics using PHYLOGIC: To evaluate the mutation clonality in the patient-matched primary and metastatic samples, we used PHYLOGIC clustering of the mutation-specific cancer cell fractions (CCFs), as previously described^13,14^. All these key steps, of the CCF and Evolutionary analysis, are found and described at http://www.broadinstitute.org/cancer/cga/acsbeta.

*Evolutionary analysis of copy-number and gene-inactivation alterations*

Corrected quantification of copy number: gene amplifications are based on the purity corrected measure for the segment containing that gene, based on ABSOLUTE^12^ (rescaled_total_cn). To better measure segment-specific copy-number, we subtracted the genome ploidy for each sample to compute copy number above ploidy (CNAP). CNAP of at least 3 are considered as amplifications (AMP), CNAP below 3 are considered low amplification and ignored in our analysis). CNAP of at least 6 are considered high amplifications (HighAMP), and CNAP of at least 9 and of segment smaller than 3000000 bp is considered very high focal amplification (FocalAMP).

For the inference of gene deletions and inactivations, we aim to infer bi-allelic inactivations (BiDel) by taking into account various events that may result in inactivation of both alleles (“two hits”). These events include: (1) loss of heterozygosity (LOH), (2) SNV (excluding the following variant classifications: "Silent", "Intron", "IGR", "5'UTR", "3'UTR", "5'Flank", "3'Flank"), (3) short indels, (4) long deletions and gene rearrangements inferred by SvABA^15^, and (5) potentially pathogenic germline events in cancer genes. Potentially pathogenic germline events include: (1) ClinVar significant annotation among the following: Pathogenic. Likely pathogenic, Conflicting interpretations of pathogenicity, risk factor; (2) Variant Classification among the following: Splice_Site, Frame_Shift_Del, Frame_Shift_Ins, Nonsense_Mutation; (3) Genome Aggregation Database (gnomAD)^16^ less than 0.05 (indicating it is a rare variant)**.**

The evolutionary classification of bi-allelic inactivation is based on comparison of the bi-allelic inactivation indicator in the pre-treatment and the post-treatment tumor samples. If BiDel was indicated for that gene in both pre and post samples, the evolutionary classification is “Shared”, if only indicated at the post-treatment sample it is “Acquired”, if only indicated at the pre-treatment sample it’s “Lost”.

The evolutionary classification of amplifications accounts for the magnitude of the observed copy-number difference between the pre-treatment and the post-treatment samples. If the difference between the CNAP of the post-treatment and the CNAP of the pre-treatment is smaller than 50%, the amplification is defined as “Shared”. If the CNAP of the post-treatment is larger than the CNAP by more than 50% and the lower pre-treatment CNAP is not at “FocalAMP” level, the evolutionary classification is “Acquired”.

If CNAP of the post-treatment is smaller by at least 50%, comparing to the pre-treatment sample and the lower post-treatment CNAP is not at “FocalAMP” level, the evolutionary classification is “Loss”. Otherwise, the evolutionary classification of amplifications is defined as “Unresolved”.

*Whole transcriptome sequencing*

For frozen tumor tissue, DNA and RNA are extracted simultaneously from a single frozen tissue or cell pellet sample using the AllPrep DNA/RNA kit (Qiagen).  For FFPE tumor tissues, DNA and RNA are extracted simultaneously using Qiagen’s AllPrep DNA/RNA FFPE kit. The resulting reads were aligned to the GRCh37 (hg19) human reference with annotations from GENCODE release 19 (GRCh37.p13) using STAR version 2.5.3a.^17,18^ Expression values were calculated using RSEM version 1.2.22.

*Fusion analysis*

Gene fusions were identified using STAR-fusion version 0.5.4 run on STAR-aligned BAM files.^19^ Candidate fusion genes were required to have the following properties: (1) A nonzero number of “JunctionReads,” reads that directly cover the fusion junction, (2) a nonzero number of “SpanningFrags,” reads where one pair aligns to the 5’ gene, and the other pair aligns to the 3’ gene, and (3) a fusion sequence that remains in-frame.

**References – Methods Section:**
