## Supplementary material for "Reversion and non-reversion mechanisms of resistance to PARP inhibitor or platinum chemotherapy in *BRCA1/2*-mutant metastatic breast cancer": Table S1 Legend and Table S2

**Supplemental Tables**

**Table S1.** *See attached Excel file.* Exact genomic coordinates, allelic fraction, and cancer cell fraction for all germline and somatic events (affecting germline-mutated gene) in *BRCA1* or *BRCA2* in the 8 patient cohort. Under biopsy ID, “A” indicates a tissue specimen obtained prior to platinum/PARPi exposure, “T” indicates a tissue specimen obtained following platinum/PARPi exposure, and “BB” indicates a blood specimen obtained following platinum/PARPi exposure. All somatic events were acquired following platinum/PARPi exposure othen than the two events indicated in gray text, which were identified prior to platinum/PARPi exposure in patient 510.

| **Patient ID** | **Number of ctDNA specimens sequenced successfully** |
| --- | --- |
| 292 | 1 |
| 303 | 3 |
| 318 | 2 |
| 339 | 2 |
| 349 | 7 |
| 359 | 2 |
| 510 | 1 |
| 565 | 0 |

**Table S2.** Number of ctDNA specimens successfully sequenced (using WES) from blood in each patient. All ctDNA specimens were drawn very close in time or subsequent to post-resistance tumor sampling.
